# Comparative pollinator conservation potential of coffee agroforestry relative to coffee monoculture and tropical rainforest in the DR Congo

**DOI:** 10.1101/2024.02.23.581744

**Authors:** Depecker Jonas, Vandelook Filip, Jordaens Kurt, Dorchin Achik, Ntumba Katshela Benjamin, Broeckhoven Ieben, Dhed’a Benoit, Devriese Arne, Deckers Lien, Stoffelen Piet, Honnay Olivier

## Abstract

Animal-pollination is crucial in the reproduction of many crops grown in the tropics, including the self-incompatible Robusta coffee. *Coffea canephora* Pierre ex A. Froehner. is indigenous to the Congo basin where it is growing in the rainforest understorey yet providing very low yields. Cultivation therefore mainly occurs in either unshaded monocultures or in agroforestry systems. Here we surveyed the Diptera (true flies) and Hymenoptera (bees) communities that are putative coffee pollinating organisms in the Yangambi region in DR Congo, and we assessed the comparative benefits of coffee agroforestry and monocultures to the coffee pollinator community. To assess a base line value of pollinator conservation value of the agroforestry system, we also compared it with natural rainforest. Using white pan traps, we identified 9,597 specimens. Natural rainforest harboured a higher number of individuals, as well as a higher number of species than both agroforestry and coffee monoculture systems, with no differences between the latter two land-uses. The Simpson diversity and Pielou’s evenness on the other hand did not differ among land-uses. Furthermore, we observed different responses in species richness and diversity to land-use between Diptera and Hymenoptera. Our analyses of pollinator community composition showed a high dissimilarity between natural rainforest and the two cultivation systems, without significant differences between the latter land-uses. Specifically, the community composition of the agroforestry and coffee monoculture systems were totally different, rather than a subset of the community composition of the natural rainforest. Our study indicates that rehabilitation of agricultural land through intercropping fruit trees may not always enhance the pollinator community and that the studied agroforestry system falls short of matching the pollinator conservation potential found in natural rainforests. A more optimal selection of tree species intercropped with coffee may both enhance the conservation value of the agroforestry system and the provisioning of pollination services.

## Introduction

Pollination is crucial for sexually reproducing plants. Worldwide, roughly 87.5 % of the flowering plant species require animal-pollination among which insects, birds, and mammals are the most prevailing pollinators (Ollerton *et al*., 2011; Osterman *et al*., 2021). Likewise, over 80 % of global crops depend, to varying extents, on animal-pollination (Klein *et al*., 2007). However, in reality, this percentage might be even higher as the true animal-pollinator dependency of crops is often underestimated (Aizen *et al*., 2009). Furthermore, agriculture in developing countries, including the tropics, is 50 % more dependent on pollinators than in developed countries, in terms of production (Aizen *et al*., 2009). Therefore, safeguarding the stability of agricultural production is key to ensure food security and livelihoods, especially in the tropics. (Driscoll *et al*., 2022). This requires detecting and conserving the crucial interactions between seed or fruit producing crops and their pollinators.

Coffee (*Coffea* spp.: Rubiaceae) is globally one of the most important agricultural commodities grown in the tropics, with an estimated annual retail market value of over 100 billion USD (Bermudez *et al*., 2022). Accordingly, the global coffee industry supports the livelihoods of over 100 million people through their involvement as smallholders, seasonal labourers, and local businesses (Guido *et al*., 2020). The coffee cultivation zone is situated between the Tropics of Cancer and Capricorn, and stretches across Africa, Central and South America, and Asia, a zone that is generally termed the coffee bean belt (Reay, 2019). Coffee production is largely dominated by *Coffea arabica* L. (Arabica coffee) (*circa* 60 %) and *Coffea canephora* Pierre ex A. Froehner (Robusta coffee) (*circa* 40 %) (ICO, 2024). The importance of *C. canephora* on the coffee market is, however, increasing because of its higher disease resistance (Leroy *et al*., 2005), higher productivity (Wellman, 1961; Campuzano-Duque *et al*., 2022), and its likely lower susceptibility to climate change than *C. arabica* (Davis *et al*., 2012; Craparo *et al*., 2015). *Coffea canephora* is native to the tropical rainforests in Africa, from Guinea in the West to Uganda in the East (Davis *et al*, 2006). Furthermore, and contrary to *C. arabica*, *C. canephora* is a self-incompatible species, making coffee yields highly dependent on robust pollinator communities (Klein *et al*., 2003a; Nowak *et al*., 2011; Bawin *et al*., 2021). Besides pollinator abundance, pollinator diversity has also been shown to be crucial in maintaining agricultural productivity of coffee (Garibaldi *et al*., 2013; Katumo *et al*., 2022). For example, the fruit set of *C. canephora* is significantly higher when flowers are visited by a species-rich, rather than a species-poor wild bee community (Munyuli, 2014). While several studies have reported that Diptera (true flies) and Hymenoptera (bees) are the main pollinators in Robusta coffee plantations (Willmer and Stone, 1989; Klein *et al*., 2003b), very little is known about the pollinator species of *C. canephora* in more natural settings, and specifically within its native range in the rainforest of the Congo Basin.

Coffee yields of wild *C. canephora* in the understorey of the tropical rainforest of DR Congo are very low because of the low population densities, and low flowering frequency and density, and possibly also the high degree of shading (Personal communication I. Broeckhoven). Cultivation of Robusta coffee therefore mainly occurs in either open, unshaded, production systems (coffee monocultures), or in agroforestry systems where coffee trees are intercropped with a selection of (fruit)trees that were re-introduced in open land, or trees that were preserved during forest clearing. Agroforests in general can either be derived through converting tropical rainforest by thinning the canopy and replacing the understorey with coffee, or through converting open land by planting trees alongside coffee, as is often the case in the DR Congo. Whereas the former is considered as forest degradation with significant loss of biodiversity and ecosystem services, the latter is considered a form of land rehabilitation with significant benefits in terms of biodiversity and ecosystem services (Klein *et al*., 2002; Martin *et al*., 2020).

In general, agroforestry practices are expected to positively affect pollinator diversity and community composition, as compared to monoculture systems (Centeno-Alvarado *et al*., 2023). This would be the case if intercropped trees provide additional nesting resources and the increased canopy cover alters the microclimatic conditions, such as light intensity and air humidity (Klein *et al*., 2002; Klein *et al*., 2003; Munyuli 2012; Centeno-Alvarado *et al*., 2023). Yet, there is a pressing need for comprehensive studies of pollinator communities across the different coffee cultivation systems in the DR Congo, where data on the composition and richness of insect communities in general are virtually lacking (Rodger *et al*., 2004; Garibaldi *et al*., 2013; Gemmill-Herren *et al*., 2014). Furthermore, although Hymenoptera have been established to be the most important coffee pollinator guild, Diptera may play a significant role in pollination as well (Orford *et al*., 2015; Rader *et al*., 2016; Doyle *et al*., 2020), although they are frequently overlooked (Ssymank *et al*., 2008). As the ecology of Diptera and Hymenoptera is quite different, these two groups are expected to respond differently to changing ecological conditions, such that Diptera may still ensure pollination services when bee populations have declined (Orford *et al*., 2015). Improving our knowledge on Diptera and Hymenoptera communities of coffee production systems and their response to agronomic practices is therefore key, especially in the Afrotropical Region.

Finally, to evaluate the role of agroforestry systems in biodiversity conservation in general, and the conservation of insect communities in particular, it is crucial to assess the (dis)similarity of the community composition and differences in species diversity between agroforestry systems on the one side and natural rainforest as benchmark on the other side (De Beenhouwer *et al*., 2013). Yet, studies directly comparing agroforestry systems of Robusta coffee with natural rainforests in terms of insect communities are rare. Based on results from studies on other tropical crops, it can be expected that a significantly different community composition of pollinators would be found between natural rainforest and agroforestry systems (Liow *et al*., 2001; Hoehn *et al*., 2010). Differences in pollinator diversity, on the other hand, are more difficult to predict as some studies have indicated an increase in bee diversity in agroforestry systems as compared to rainforest (Bos *et al*., 2007; Hoehn *et al*., 2010).

In this study, we used 600 white pan traps across 30 plots to survey the Hymenoptera and Diptera communities as the putative main coffee pollinators in the Yangambi region in DR Congo. We aimed to assess to what extent agroforestry practices of Robusta coffee benefit the coffee pollinator diversity and differ from the community composition found in coffee monocultures. Furthermore, to assess the conservation value of the agroforestry system, we aimed to compare the pollinator community composition and diversity of the agroforestry system to those in natural rainforest sites as a benchmark. Finally, we aimed to assess whether Hymenoptera and Diptera communities responded differently to land-use. Overall, we hypothesised to find increasingly impoverished pollinator communities along the gradient from natural rainforest over agroforestry to coffee monoculture, with the Diptera and Hymenoptera communities of agroforests and coffee monocultures being a subset of the communities in the natural rainforest.

## Material and methods

### Study area and survey plots

Sampling occurred between January 2022 and March 2022 in the Yangambi region, in the Tshopo province, in the North-Eastern part of the Democratic Republic of the Congo, approx. 100 km west of Kisangani. The prevailing climate can be characterised by two drier seasons (December–March and June–July) and two rainy seasons (April–May and August– November) (van Vliet *et al*., 2018). The landscape of Yangambi is a typical forested ecosystem of the Congo Basin and consists of a mosaic of land tenures: the Yangambi Man and Biosphere Reserve, the Ngazi Forest Reserve, a logging concession, and customary land (van Vliet *et al*., 2018). For this study, a total of 30 survey plots of 25 m x 25 m were established in three different land-use systems during the main coffee flowering period (January–March). Ten plots were established in the natural rainforest (FOR), ten plots in monocultures of *C. canephora* (MONO), and ten plots in *C. canephora* agroforests (AGRO). The plots in natural rainforests were characterised by a high level of canopy closure (91 % ± 1 %), with a diverse mixture of tree species such as *Cola griseiflora* De Wild. and *Staudtia kamerunensis* Warb. and with a small number of *C. canephora* individuals. In the coffee monocultures, a tree canopy was entirely absent as *C. canephora* is the only arboreal species present. In the agroforestry system, the canopy displays medium levels of closure (45 % ± 3%), and the arboreal vegetation consists of a mixture of planted fruit trees like *Pachylobus edulis* G.Don., *Elaeis guineensis* Jacq., and *Persea americana* Mill., and very rarely also native rainforest trees like *Anonidium mannii* (Oliv.) Engl. & Diels. *and Barteria nigritana* Hook. f..

### Pollinator sampling and identification

In each of the 30 survey plots, five white pan traps were placed in an approximately x-shaped design with an inter-distance of at least 5 m following Droege *et al*. (2010). The pan traps were 130 mm in diameter and 60 mm deep. Pan traps were three quarters filled with water and a few drops of detergent were added to break the surface tension to prevent insects from escaping while in the water. The pan traps were placed on a wooden stick at breast height and were active for four weeks and emptied at regular time intervals. Loss of 10 % of the pan traps resulted in their shorter operation time of only three weeks. Insects were preserved in 70% ethanol for later identification. Samples were transported to the RMCA where they were sorted to insect order. Then, all Hymenoptera (bees) and Diptera (true flies) were identified by expert taxonomists to genus or species level, where possible (see in the Acknowledgments). For certain groups, for which expert taxonomists were not available, we identified the specimens to morphospecies, an appropriate surrogate for formal species, commonly used in ecological studies (Oliver and Beattie, 1996). In the remainder of the text, we will refer to “putative pollinators” as to “pollinators” for simplicity, without confirming the actual contribution of the species to pollination.

### Data analysis

All statistical analyses were performed using R software (R Core Team, 2023), including combined and separate analyses of Diptera and Hymenoptera combined. In all analyses, the data of the five pan traps were pooled for each survey plot. Prior to further data analyses, single- and doubletons were removed from the dataset, keeping only taxa with at least three individuals across all survey plots (see Coddington *et al*., 2009; Barlow *et al*., 2010; Allen *et al*., 2016).

For each survey plot, species richness (Hill’s N0), species diversity (Hill’s N1 or the exponential of the Shannon entropy, and Hill’s N2 or inverse Simpson index), and Pielou’s evenness (Hill’s N1/log(Hill’s N0)) were calculated based on the collected species abundance data with 9,999 bootstraps using the iNEXT function of the iNEXT package (Hsieh *et al*., 2016). These estimates were afterwards standardised for sample completeness, as sample sizes differed between survey plots. Here, sample completeness represents the proportion of the total number of individuals in a unit that belongs to the species in the sample. We opted to standardise for sample completeness rather than sample size, as the former allows less biased comparisons (Chao and Jost, 2012). Following Chao and Jost (2012), the extrapolation of the estimates was done to double the reference sample size per survey plot. Potential differences in abundance, species richness, species diversity, and Pielou’s evenness between the three land-use systems were analysed by using an Analysis of Variance (ANOVA). All metrices followed the assumptions for parametric testing, except abundance, which was log-transformed to comply with the assumptions for parametric testing. Tukey’s Honestly Significant Differences (HSD) tests were used to explore significant outcomes of the ANOVA tests further. For this, abundance and Hill’s N1 were log-transformed for the Diptera, while abundance, species richness, Hill’s N2, and Pielou’s evenness, were log-transformed for Hymenoptera.

Comparison of the pollinator community composition across the three land-uses was done in two ways. First, using the *metaMDS* function in the vegan package (Oksanen *et al*., 2020), the dissimilarity in community composition was visualised using non-metric multidimensional scaling (NMDS), based on the Bray-Curtis dissimilarity measure at survey plot level, which is the most suitable dissimilarity measure for abundance data (Ricotta and Podani, 2017). The provided two-axis solution was a good representation of the data in reduced dimensions, as indicated by the stress level (0.11) which was well below the 0.20 threshold (Kruskal, 1964). Second, using the *adonis* function in the Vegan package (Oksanen *et al*., 2020), where differences in the community composition between the three land-uses were tested by a permutational multivariate analysis of variance with 9,999 permutations. Subsequently, the observed differences were analysed in more detail through pairwise comparisons of the different land-uses with 9,999 permutations with Bonferroni correction, using the *pairwise.adonis* function in the pairwiseAdonis package (Arbizu, 2017). Additionally, the effect of geographic distance on the dissimilarity of the pollinator community composition was tested by performing a Mantel test with 9,999 permutations using the *mantel* function in the Vegan package (Oksanen *et al*., 2020). The Bray-Curtis dissimilarity measure was also calculated at land-use level, with survey plots pooled per land-use. Across the three different land-uses, indicative pollinator taxa were identified based on their indicator value by a multilevel pattern analysis using the *multipatt* function in the *Indicspecies* package (Dufrêne and Legendre, 1997; De Cáceres and Legendre, 2009).

Nestedness of the pollinator communities between the three land-uses, *i.e.* whether the pollinator communities of agroforest sites and coffee monoculture were a subset of the communities found in the rainforests, was assessed through the nestedness metric based on overlap and decreasing fill (NODF index) (Almeida-Neto *et al*., 2008) and validated against null matrices of a null model. Prior to calculating the NODF, insect abundances were transformed to a presence-absence matrix (1=present, 0=absent), in which columns and rows represented species and land-use systems, respectively, and with survey plots pooled per land-use system. Instead of arranging the rows by the marginal row totals, we opted to arrange the rows by the different land-uses, to allow to test for the prior specified hypotheses. The *nestednodf* function in the Vegan package (Oksanen *et al*, 2020) was then applied using values of the resulting matrix to calculate the NODF. This nestedness metric described varies between 0 (not nested) and 100 (perfectly nested) and is considered more appropriate than other nestedness meteics as NODF is quantified for rows and columns independently, allowing separate evaluation of the nestedness among systems and species, respectively (Almeida-Neto *et al*., 2008; Sasaki *et al*., 2012). Finally, the significance of the calculations was evaluated against the “curveball” null model (Strona *et al*., 2014) using the *oecosimu* function in the Vegan package (Oksanen *et al*., 2020) with 100,000 simulated null communities and 5,000 simulations burned. The “curveball” method generates null matrices with the same row and column marginal totals as the original presence-absence matrix, which helps minimise the risk of type-II errors to conserve ecologically relevant information (Joppa *et al*., 2010; Strona *et al*., 2014), and is therefore advantageous over other null models.

## Results

With the removal of 108 singletons and 49 doubletons (see appendix), a total of 9,389 Diptera and Hymenoptera specimens were sampled in the 30 survey plots across three different land-uses over a period of four weeks. Within the Diptera, the 8,542 individuals belonged to 149 taxa, of which 88 % were identified to (morpho)species level, 8 % to genus level, and 4 % were unidentified (most of these because of damage during the processing and the transport of samples). Within the Hymenoptera, 847 individuals belonged to 31 taxa, of which all were identified to (morpho)species level. Numbers of shared and unique taxa across the different land-use systems are displayed in Fig. 2.

**Figure 1:**
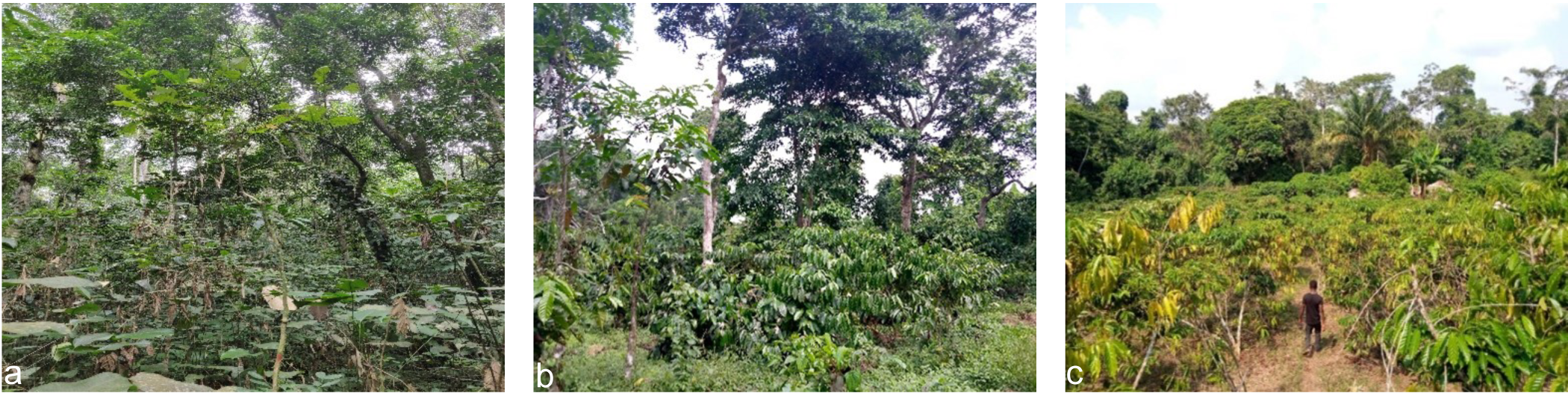
Overview of the three land-use systems in the Yangambi region in the DR Congo. (a) Natural rainforest (FOR), (b) Agroforestry (AGRO), and (c) coffee monoculture (MONO). © Ieben Broeckhoven

**Figure 2:**
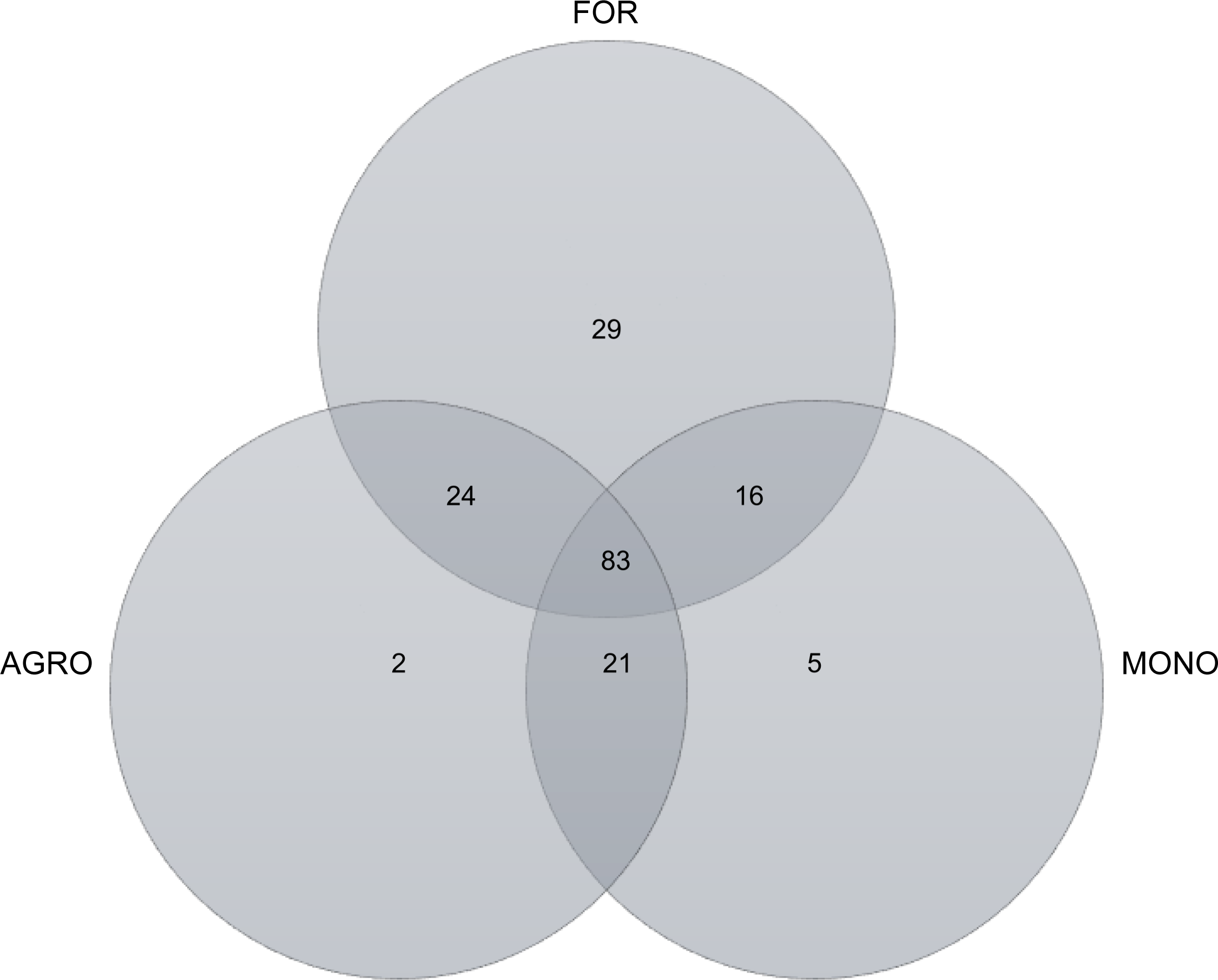
Venn diagram depicting shared and unique Diptera and Hymenoptera taxa across the three land-use systems in the Yangambi region in the DR Congo.

### Species richness and diversity

Across all samples, the ANOVA tests indicated no significant differences between Hill’s N2 (F = 1.70; P = 0.201) and Pielou’s evenness (F = 1.37; P = 0.270) across the three land-uses, whereas abundance (F = 10.75; P < 0.001), species richness (F = 9.89; P < 0.001), and Hill’s N1 (F = 3.85; P = 0.034) differed among the different land-uses. Specifically, both agroforest sites and coffee monocultures yielded 73 % (P = 0.002) and 76 % (P < 0.001) fewer individuals, respectively, as compared to the natural rainforest sites. In terms of species richness, both agroforest sites and coffee monocultures harboured 26 % (P = 0.017) and 38 % (P < 0.001) fewer species, respectively, than the natural rainforest sites. In terms of species diversity (Hill’s N1), only coffee monocultures were significantly less diverse than the natural rainforest sites, with a 22 % (P = 0.026) difference (Fig. 3).

**Figure 3:**
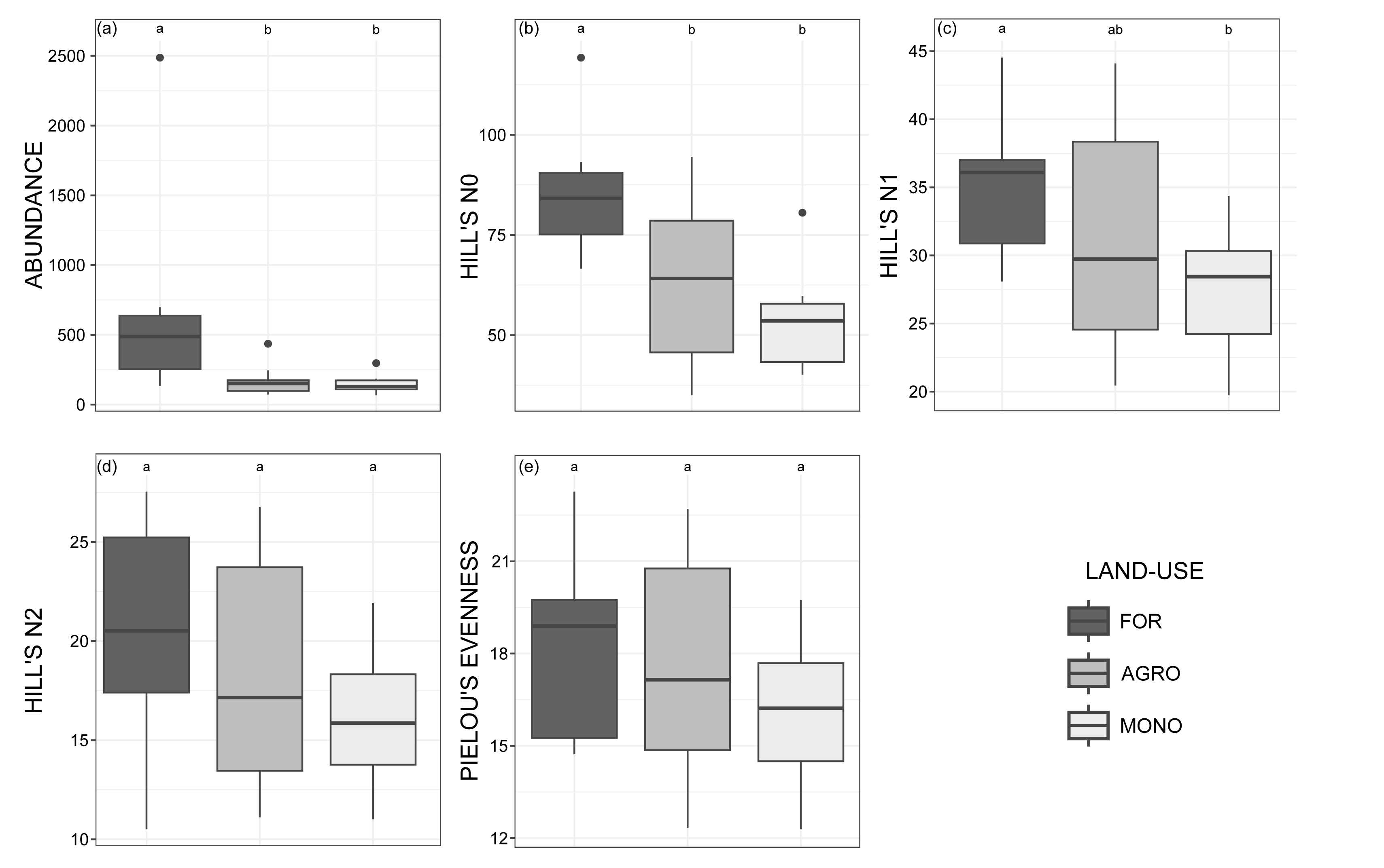
Comparisons between Diptera and Hymenoptera abundance and species diversity metrics across natural rainforest (FOR), agroforestry (AGOR), and coffee monoculture (MONO) in the Yangambi region (DR Congo). Hinges represent the 25^th^, 50^th^, and 75^th^ percentiles, respectively. Whiskers extend to maximum 1.5 times the interquartile range. Letters code for significant differences among land-uses. (a) abundance, (b) species richness, (c) species diversity 1, (d) species diversity 2, (e) Pielou’s evenness

When only considering Diptera, significant differences in abundance (F = 10.36; P < 0.001), species richness (F = 11.69; P < 0.001), and Hill’s N1 (F = 5.04; P = 0.014) were found among the three land-uses. Both agroforest sites and coffee monocultures yielded 71 % (P = 0.002) and 76 % (P < 0.001) less Diptera individuals, respectively, than the natural rainforest sites. Likewise, these land-uses were also less species rich, with 23 % (P = 0.015) and 37 % (P < 0.001) less Diptera species in agroforest sites and coffee monocultures, respectively, as compared to the natural rainforest sites. Hill’s N1 on the other hand, was only significantly lower in coffee monocultures than in the natural rainforest sites (P = 0.010), with a 23 % difference in diversity. When considering only Hymenoptera, only abundance differed significantly among the three land-uses (F = 4.90; P = 0.015). Particularly, in both agroforest sites (P = 0.033) and coffee monocultures (P = 0.027) significantly less Hymenoptera were observed (84 % and 82 % less, respectively) as compared to the natural rainforest sites (Fig. S1 and S2).

### Community composition

The ordination of the 30 survey plots showed a clear pattern of separation of the survey plots according to land-use for all taxa (Fig. 4), which was supported by the permutational multivariate analysis of variance, (R^2^ = 4.67; P < 0.001). Specifically, significant differences were found between the community composition of the natural rainforest and agroforest sites (R^2^ = 0.24; P < 0.001), and between the natural rainforest sites and coffee monocultures (R^2^ = 0.28; P < 0.001). No differences were found between agroforest sites and coffee monocultures (R^2^ = 0.06; P = 0.67). Similar results were found for both the Diptera and Hymenoptera separately (see Fig. S2). The Bray-Curtis measure showed a high dissimilarity between both the natural rainforest and agroforest sites (0.66), and between the natural rainforest sites and coffee monocultures (0.72), and a lower dissimilarity of 0.27 between agroforest sites and coffee monocultures. The Mantel test indicated no significant correlation between the Bray-Curtis dissimilarity and geographic distance between the survey plots (R = −0.02; P = 0.546).

**Figure 4:**
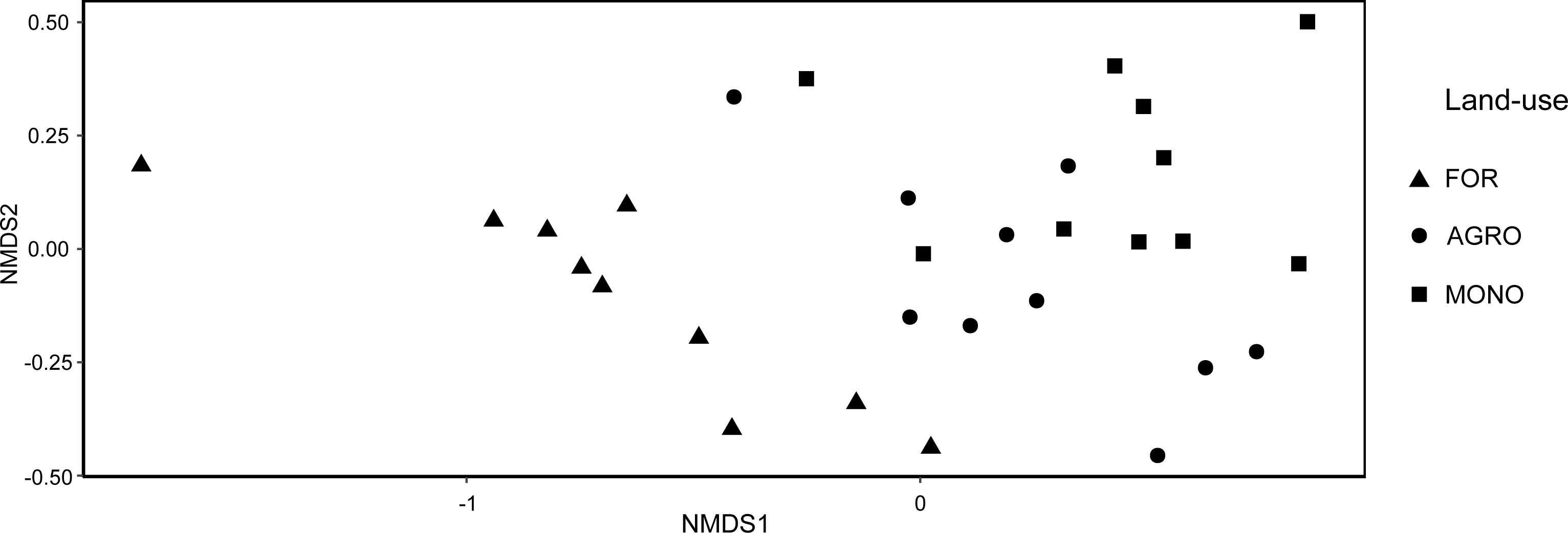
Non-metric multidimensional scaling (NMDS) ordination of 30 survey plots in the Yangambi region (DR Congo) across the three land-use systems: natural rainforest (FOR), agroforest (AGRO), and coffee monoculture (MONO). Ordination based on the Bray-Curtis dissimilarity measure calculated from the Diptera and Hymenoptera abundances and represent their community composition.

The multilevel pattern analysis revealed that 57 taxa out of the 180 were indicative across the 30 survey plots based on their indicator value (both P < 0.05 and indicator value > 0.600). A total of 50 taxa were found to be indicators for the natural rainforest sites (41 Diptera and 9 Hymenoptera), one Diptera for agroforest sites, and six for coffee monocultures (5 Diptera and 1 Hymenoptera) (Table S1). Results of the multilevel pattern analysis of both Diptera and Hymenoptera can be found in the supplementary material.

### Nestedness

The total NODF of the presence-absence matrix of all taxa revealed a significant nested pattern (NODF = 58.88; P < 0.001). More specifically, significant nestedness was found in the columns (species) (Ncolumns = 58.88; P < 0.001), but not in the rows (land-uses) (Nrows = 58.83; P = 0.25) (Fig. 5). The community composition of the agroforest sites and coffee monocultures thus do not consists of a subset of the community composition of the natural rainforest sites. Similar patterns of nestedness were found in the analysis based on the presence-absence matrix of the Diptera and Hymenoptera datasets separately.

**Figure 5:**
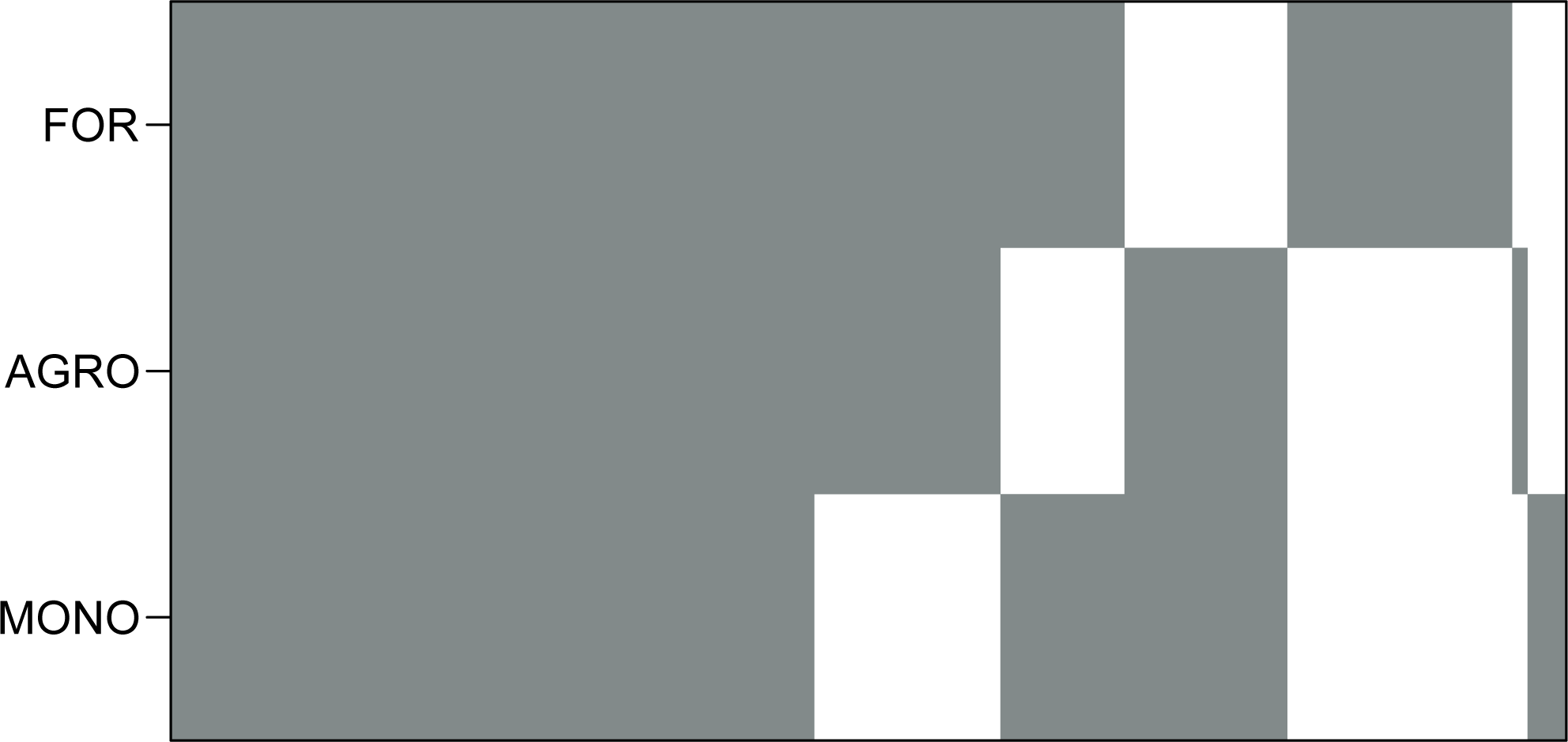
Incidence plot of Diptera and Hymenoptera species across the three land-use systems in the Yangambi region (DR Congo): natural rainforest (FOR), agroforestry (AGRO), and coffee monoculture (MONO), and on which the nestedness analysis was based.

## Discussion

An improved understanding of how agroforestry systems may benefit coffee pollinators and how they compare to the pollinator communities in undisturbed rainforest is key for both agricultural production and biodiversity conservation. Especially in the DR Congo such data are extremely scarce. Here, we compared species diversity and community composition of Diptera and Hymenoptera, the likely main coffee pollinators, between *C. canephora* agroforestry system, *C. canephora* monocultures, and natural tropical rainforests where *C. canephora* is naturally occurring in the understorey.

### Mitigating effects of agroforestry as compared to monocultures

In contrast to what we hypothesised, significant differences between the coffee monocultures and the coffee agroforest sites were neither observed in terms of abundance, species richness and species diversity, nor in terms of community composition of each the Diptera or Hymenoptera. Previous studies suggested that higher light intensity, which is strongly related to canopy cover, increased insect richness, specifically bee species richness (Klein et al., 2003b; Klein et al., 2003c; Munyuli *et* al., 2012). Furthermore, studies on *C. arabica* flower-visiting insects in Ethiopian moist evergreen forests, found a decreased species diversity with increasing agricultural and forest management intensification (Berecha *et al*., 2015; Geeraert *et al*., 2019). A possible explanation for our observations could be related to the phenology of the intercropped trees. A phenological mismatch between the flowering time of coffee and that of the intercropped trees would suggest that the trees fail to augment floral resources during the main coffee flowering period, a key driver in attracting pollinators (Meyer *et al*., 2009; Roulston and Goodell, 2011; Berecha *et al*., 2015; Krishnan *et al*., 2018). Additionally, the intercropped trees may not always provide sufficient or suitable nesting sites. Different tree species exhibit differences in wood composition and growth patterns, influencing their tendency to develop cavities, which serve as preferred nesting sites for certain pollinators (Ricketts, 2004; Roulston and Goodell, 2011). As such, nests of certain bee species (e.g. stingless honey bees of the tribe Meliponini) were found to be more associated with very large tropical rainforest trees (Eltz *et al*., 2003), which are less represented in the agroforestry systems. Furthermore, the surrounding tropical rainforest matrix may have a strongly mitigating influence on the pollinator diversity and community (Ricketts *et al*., 2008; Klein 2009; Geeraert *et al*., 2020). Given the proximity of both land-uses to the rainforest in Yangambi, with some being less than 100 meters away and the furthest just 1.4 km, the surrounding tropical rainforest might provide ample nesting resources and forage space in close proximity to the cultivation systems. We found no significant correlation between species richness and diversity, in agroforest sites and coffee monocultures and distance to the nearest rainforest margin (results not shown). In this context, the introduction of the current fruit trees may not significantly enhance pollination benefits. Interestingly, in the same survey plots, no significant difference was observed between agroforest sites intercropped with fruit trees and coffee monoculture in terms of coffee yield per plant. Whereas in the agroforest sites the mean yield was 0.96 kg of green coffee per plant, the mean yield in the coffee monoculture was 0.92 kg of green coffee per plant (I. Broeckhoven, unpublished results). The uniformity in yield per plant between the two land-uses might be associated to the absence of differences in species diversity in pollinators we found in our study (Klein et al., 2003c; Garibaldi et al., 2013; Katumo et al., 2022).

### Agroforests cannot replace rainforest in terms of conservation value

In order to assess the conservation potential of the agroforestry system, Diptera and Hymenoptera diversity and composition of the agroforestry system were compared with that of the natural rainforest. In line with our hypothesis, abundance, and species richness were significantly higher in the natural rainforest as compared to the agroforestry system. Furthermore, more single- and doubletons were removed from the natural rainforest than from the agroforestry system. This seems to indicate that the rare species are lost, while the typical and dominant species seem to prevail in the explored agroforestry system (Chao *et al*., 2014). It is broadly acknowledged that rare species tend to be more sensitive to disturbance, and show a faster loss under land-use than common species (Davies *et al*., 2004; Sekercioglu *et al*., 2008; Leitão *et al*., 2016). For example, a study on bee species in a temperate zone found less than half of rare bee species in agricultural landscapes as compared to natural forest (Harrison *et al*., 2019). In the Afrotropical Region, however, more studies are needed to verify whether rare species disappear at a faster rate than more common species with agricultural intensification. Although significant species loss could be noticed as compared to natural rainforest, agroforestry systems still tend to harbour a relatively large diversity of pollinators, which is confirmed by our nonsignificant lower estimates of species diversities in the agroforestry system. Our results are in line with those of studies, mainly on bees, in Ecuador and Indonesia that did not detect changes, or even observed higher diversities in agroforests than in the natural forests (Klein et al., 2002; Bos et al. 2007; Tscharntke et al., 2008; Hoehn et al., 2010).

Yet, as already indicated by studies on tropical tree species (Edwards *et al*., 2014; Imai *et al*., 2014; Depecker *et al*., 2022) and reported by Akter *et al*. (2020) for pollinators in a mangrove forest in Bangladesh, the impact of changes is better reflected by species community composition than by species diversity. Indeed, we observed a clear difference in pollinator community composition between natural rainforests, the agroforestry system, and the coffee monoculture system. This trend does not seem to be driven by geographic distance, as indicated by the Mantel test. Moreover, the nestedness analysis revealed that the pollinator community composition in the agroforestry system is not a subset of the community composition of the natural rainforest, but rather represent a unique assemblage of species. This might be attributed to the nature of the considered agroforest system, which in Yangambi represents a rehabilitated fallow land rather than a result of the conversion of tropical rainforest into agroforest by forest thinning (Martin *et al*., 2020). Similarly, pollinator communities in coffee agroforests were found to be significantly different from those in more natural habitats in Ethiopia and Indonesia (Klein *et al*., 2002; Hoehn *et al*., 2010; Berecha *et al*., 2015).

Despite the clear distinctions between different systems in terms of community composition, it is important to note that there were still shared taxa among land-use systems. In fact, a substantial number of taxa is shared, although their abundances vary considerably. For example, 323 individuals of *Meliponula bocandei* Spinola. were recorded in the rainforest, whereas only one individual was found in the coffee monoculture system. Because this species typically nests in cavities of rainforest trees like *Albizia gummifera* (J.F.Gmel.) C.A.Sm. (Pauly and Vereecken, 2013; GBIF Secretariat, 2023) that are absent in the agroforestry system, the absence of this species in this system is no surprise. Likewise, while 93 *Graptomyza triangulifera* Bigot. individuals were observed in the rainforest, only one was observed in the agroforestry system, although it is considered a generalist pollen collector (ICIPE, 2023). Yet, crucial ecological information for this species is still lacking, as is for the majority of the Diptera, to clearly understand the observed patterns. Whereas autecological information is lacking for many species, at the family level broad trends can be observed. Many Drosophilidae and Platystomatidae prefer shaded and densely vegetated areas, and are therefore likely more recorded in the natural rainforest, whereas Chloropidae are abundant across diverse habitats, potentially accounting for their equally abundant presence in all three land-use systems (Kirk-Spriggs and Sinclair, 2017). A species much more abundant in both the agroforest sites and coffee monocultures, as compared to natural rainforest sites, was *Trirhithrum coffeae* Bezzi, which is the main fruit fly pests of coffee (Agricultural Research Council, 2023). Overall, our results suggest rainforest specialist species are not preserved in the agroforestry system, underscoring the irreplaceable role of tropical rainforests in biodiversity conservation (De Beenhouwer *et al*., 2013).

### Bees respond differently than flies to changes in land-use

Nearly all studies comparing pollinator diversity and composition between coffee cultivation systems and tropical rainforest only focus on Hymenoptera, despite the recognised importance of Diptera in pollination (Orford *et al*., 2015; Rader *et al*., 2016; Doyle *et al*., 2020). Our findings highlight the importance of considering Diptera as well, as they respond differently to land-use. Diptera differed in terms of abundance, species richness, and species diversity (Hill’s N1) between land-use systems, whereas for Hymenoptera only abundance differed between land-use systems. In terms of community composition, however, similar patterns were observed in Diptera and Hymenoptera. Similarly to our study, Geeraert *et al*. (2019) compared semi-natural coffee forests and shaded coffee plantations in the highlands of SW Ethiopia and found differences in non-*Apis* bee and hoverfly abundance, but not in the abundance of other Diptera. The differences we observed might be associated with the ecology of the different taxa. Tropical bee communities include more social species that are floral generalists by definition as compared to more arid regions (Michener, 2007; Wood *et al*., 2023) and are often more constrained by nesting resources, whereas flies have much broader dietary requirements (Staton *et al*., 2022). However, additional research is needed to elucidate the differences in responses to agricultural practices, encompassing both taxonomic and ecological aspects. In the realm of taxonomy and ecology of tropical pollinators, much work remains to be done, including investigations into functional traits.

## Conclusion

We found differences in both the Diptera and Hymenoptera community composition among various land-uses in the Yangambi area in the DRC with rainforest harbouring by far the highest diversity showing its irreplaceable conservation value. Despite the common assumption that the introduction of trees in agricultural land contributes positively to biodiversity our study revealed no discernible impact on the coffee pollinator community when intercropping coffee monocultures with trees. We suggest that this is due to both characteristics of the introduced trees and the presence of sufficient suitable habitat in the nearby rainforest. Nevertheless, additional research is essential to examine the possibility of a delayed response, encompassing both buffering and depleting effects, in terms of biodiversity and pollination services. Furthermore, the predominant intercropped species in the examined agroforestry system are fruit trees and further research may assess the impact on pollinators when intercropping with trees that are native to the local rainforest. Furthermore, it is important to note that our sampling approach did not allow to characterise flower-visiting insects and that the functional pollinator community of *C. canephora* in Yangambi remains unexplored.

## Acknowledgments

First, we are grateful to all the experts who have helped with identifying the specimens: Burgert Muller for the Anthomyiidae, Faniidae, and Muscidae; Arianna Thomas for the Calliphoridae; Hans Feijen for the Diopsidae; Iain McGowan for the Lonchaeidae; Daniel Whitmore for the Sarcophagidae; Martin Hauser for the Stratiomyidae; Marc De Meyer for the Tephritidae. Syrphidae were identified by KJ, and Hymenoptera by AD. Further we would like to thank the Institut National pour l’Etude et la Recherche Agronomiques (INERA) and the FORETS project, which is financed by the 11^th^ European Development Fund, for facilitating the field mission. Finally, we would like to thank the Ministère de L’Environnement et Développement Durables (MEDD) for their help with obtaining permits (N°001/ANCCB-RDC/SG-EDD/BTB/01/2022).

## Author contributions

JD, OH, FV, and KJ designed this study. JD and BN collected the samples in the field. PS and FV helped in obtaining the permits. JD, KJ, LD, and AD identified the specimens. JD analysed the data. JD, KJ, FV, OH wrote the manuscript. All authors contributed to finalising the manuscript.

## Data availability statement

The data that support the findings of this study will be made openly available in GBIF.

## Supplementary material

Table S1: List of all encountered taxa.

Table S2: Indicator species across the three land-use systems in the Yangambi region in DR Congo.

Figure S1: Comparisons between Hymenoptera abundance and species diversity metrics across natural rainforest (FOR), agroforestry (AGOR), and coffee monoculture (MONO) in the Yangambi region (DR Congo). Hinges represent the 25th, 50th, and 75th percentiles, respectively. Whiskers extend to maximum 1.5 times the interquartile range. Letters code for significant differences among land-uses. (a) abundance, (b) species richness, (c) species diversity 1, (d) species diversity 2, (e) Pielou’s evenness

Figure S2: Comparisons between Diptera abundance and species diversity metrics across natural rainforest (FOR), agroforestry (AGOR), and coffee monoculture (MONO) in the Yangambi region (DR Congo). Hinges represent the 25^th^, 50^th^, and 75^th^ percentiles, respectively. Whiskers extend to maximum 1.5 times the interquartile range. Letters code for significant differences among land-uses. (a) abundance, (b) species richness, (c) species diversity 1, (d) species diversity 2, (e) Pielou’s evenness

