## Supplementary figures and images for "Comparative pollinator conservation potential of coffee agroforestry relative to coffee monoculture and tropical rainforest in the DR Congo"

### Supplemental figure S1

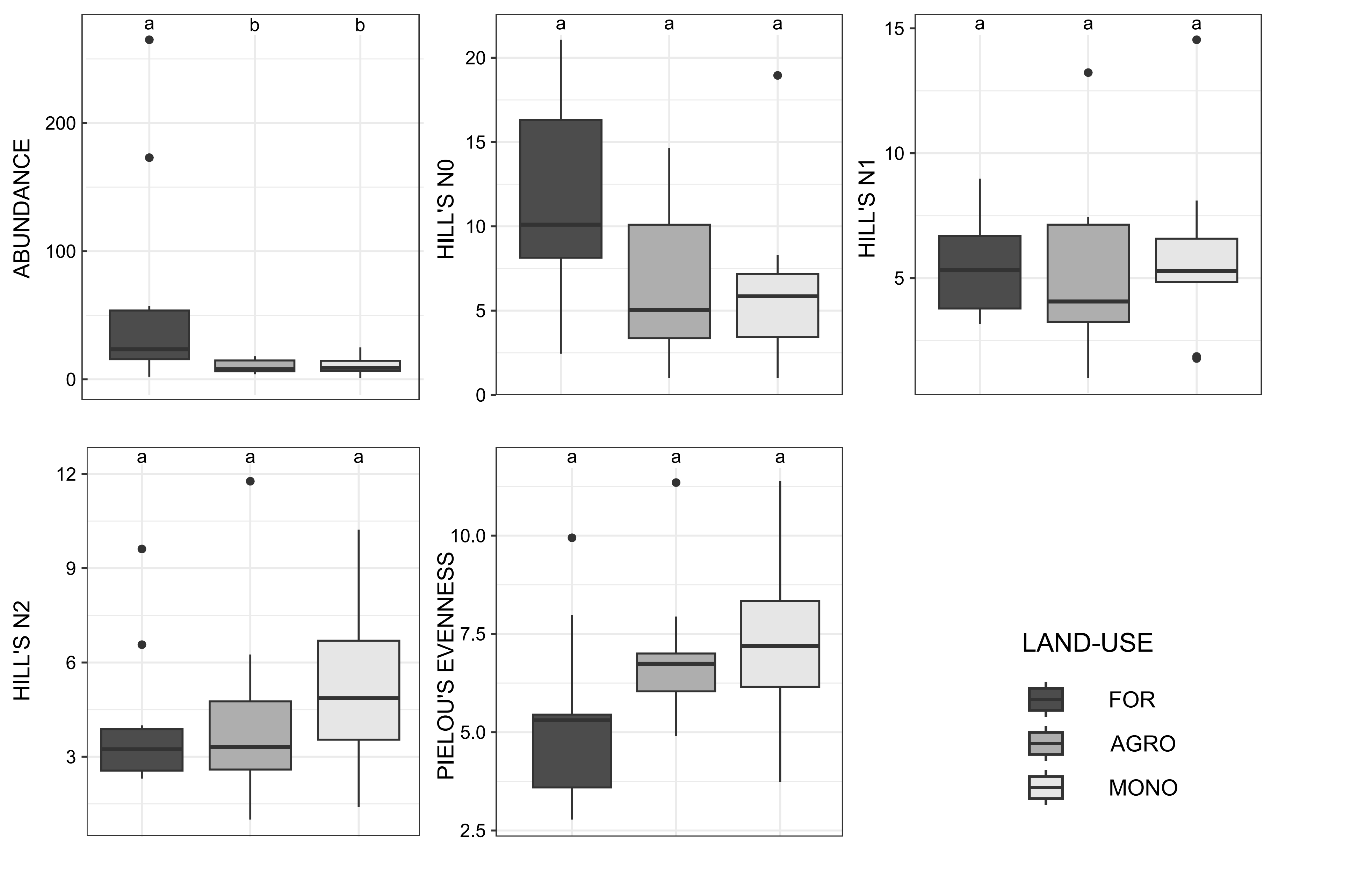

### Supplemental figure S2

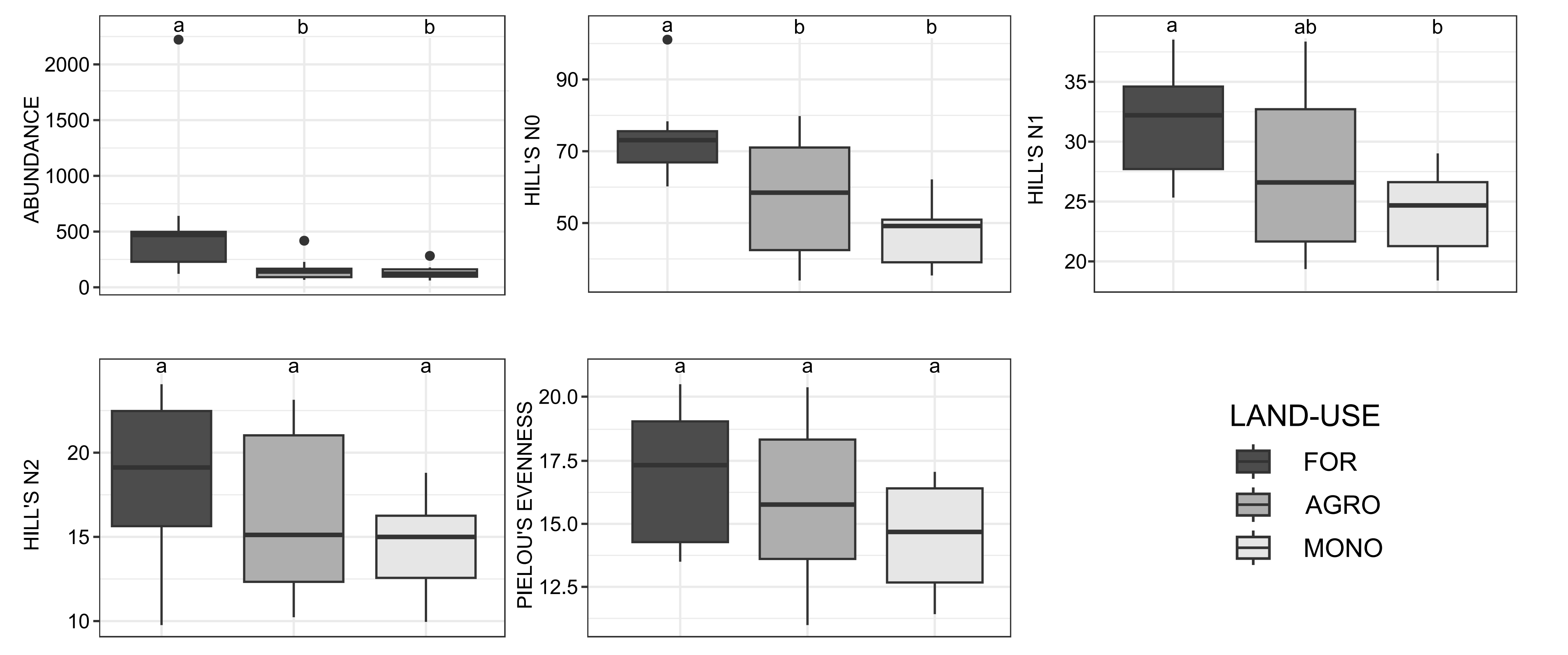
